# Assessing the impact of spatial and temporal filters on BirdNET performance for monitoring bird communities

**DOI:** 10.64898/2026.06.02.729608

**Authors:** Cristian Pérez-Granados, Jon Morant, David Funosas, Esther Sebastián-González

## Abstract

Recent advances in automated technologies, such as passive acoustic monitoring, provide a powerful framework for surveying bird communities at broad spatial scales. Among the most widely used artificial intelligence tools for automated bird sound recognition is BirdNET, which can identify over 6,000 species worldwide. However, the effects of key user-defined settings, such as species filtering, remain poorly evaluated. Here, we assess how alternative species-filtering strategies influence BirdNET performance in describing bird communities worldwide. We analysed 5,047 minutes of sound recordings from 72 locations worldwide, comprising 1,192 bird species identified by expert ornithologists. We compared three common species-filtering approaches applied in BirdNET workflows to post-process its output: no filtering, spatial filtering (species present all-year at a given location), and spatio-temporal filtering (species present at a given location and week). The unfiltered approach maximised BirdNET species detection (recall) but suffered very low precision (had many misidentifications) and poor overall performance. In contrast, the other two filtering strategies greatly improved precision and overall performance, despite moderate reductions in recall. Among them, spatio-temporal filtering consistently achieved the best performance across most datasets and regions globally. Within this optimal filtering approach, we also evaluated the role of another parameter: occurrence probability thresholds. Intermediate values of this threshold (around 0.05) maximized BirdNET performance in community-level analyses. Our results demonstrate that species filtering is a key but often underappreciated component of BirdNET workflows. We hope our findings may guide future studies in selecting optimal species filters, while emphasising that filtering selection should be guided by study objectives and data context.

## INTRODUCTION

Automated and non-invasive approaches have become increasingly important in biodiversity monitoring (Lahoz-Monfort and Magrath 2021). Among these, passive acoustic monitoring (hereafter PAM) has proven particularly effective for surveying a wide range of taxa (Sugai et al. 2019, Darras et al. 2025). Birds represent the group most frequently monitored with PAM (Shonfield and Bayne 2017, Sugai et al. 2019), as it enables a reliable characterization of bird communities (Darras et al. 2018) and the estimation of population densities from passively recorded soundscapes (Pérez-Granados and Traba 2021). Recent improvements in automated signal recognition, particularly deep learning models, have further expanded PAM application for avian monitoring (Stowell 2022, Xie et al. 2022). Taking advantage of these methodological advances, several user-friendly bird recognition tools have been introduced in recent years, streamlining PAM workflows and making them more accessible to practitioners, the general public and researchers with limited informatics background. Among these, BirdNET is particularly notable because of its broad taxonomic and geographic scope, as well as its potentially high predictive accuracy (Kahl et al. 2021; Funosas et al. 2024, 2026).

BirdNET is a free, user-friendly bird recognition software based on a convolutional neural network (Kahl et al. 2021). The latest BirdNET model (v.2.4) can identify the vocalizations of more than 6,000 bird species, which are recognized in 3-second windows. Recent research has proven both its potential and limitations for detecting bird vocalizations and describing bird communities worldwide (Pérez-Granados 2023, Funosas et al. 2024, 2026). BirdNET has three customizable inference settings with a high impact on its output: *Minimum confidence score threshold, Overlap*, and *Sensitivity*. Each BirdNET prediction has an associated confidence score, which ranges from 0.01 (very low model certainty in the prediction) to 1 (very high certainty). Users can filter BirdNET output by setting a customizable *Minimum confidence* score threshold. Setting a low threshold increases the probability of detecting a bird vocalization in the dataset but also the risk of false positives, while setting a high threshold reduces errors at the cost of losing some correct predictions (Wood and Kahl 2024). The *Overlap* (ranging from 0 to 3 seconds) parameter determines the degree of temporal overlap between consecutive 3-second windows, whereas *Sensitivity* (ranging from 0.5 to 1.5) adjusts how confidence scores are distributed across predictions (see extended descriptions of the three customizable settings in Pérez-Granados et al. 2026a). A recent study conducted on a global scale assessed the impact of these three settings on BirdNET performance and suggested that the optimal configuration of these parameters differs among species, regions, and research goals (Pérez-Granados et al. 2026a).

In addition to these settings, BirdNET allows users to restrict the species included in the output by applying spatial and spatio-temporal filters. When no filter is applied, BirdNET scans the recordings using the full list of acoustic categories, including abiotic sound sources, on which the model was trained (Kahl et al. 2021). This approach increases the probability of detecting rare or vagrant species in the datasets, but often leads to multiple predictions of species that do not occur in the monitored area. Therefore, a common approach is to upload a custom species list to constrain BirdNET output to a predefined set of target species (Cole et al. 2022, Manzano-Rubio et al. 2022; Bota et al. 2024). Users can use lists automatically generated by BirdNET by providing the geographic location and date of the recording. The lists can include: 1) all the species predicted to occur in that area at any time of the year (“*Spatial filte*r”) or 2) only those species predicted to occur in that area during that specific week of the year, by also specifying the recording week (“*Spatio-temporal filte*r”). Moreover, automatic BirdNET lists are filtered to include only those species occurring above a user-defined minimum occurrence frequency threshold (hereinafter referred to as probability of occurrence). The probability of occurrence setting ranges from 0.01 to 0.99 (default 0.03) and it is estimated using eBird checklist frequency data for a certain latitude, longitude, and week of the year. Setting a low probability of occurrence results in broader species lists (i.e., including species with low predicted occurrence probabilities), whereas higher thresholds restrict the lists to species with higher expected occurrence (i.e., excluding those with lower probability of occurrence).

eBird is a global, citizen-science repository of bird observations (Sullivan et al. 2014), whose data are known to exhibit spatial biases in sampling effort. In fact, according to BirdNET developers, such occurrence models perform reliably in regions with high eBird data coverage, such as North and South America, Europe, India, and Australia. In contrast, in regions with limited eBird observations, the generated lists may poorly reflect true occurrence probabilities, but can be used as binary filters by indicating whether a species may occur or not (see https://github.com/birdnet-team/BirdNET-Analyzer/discussions/234). Previous studies conducted on a worldwide scale have adopted a fixed probability of occurrence of 0.02 (Funosas et al. 2024, 2026, Pérez-Granados et al. 2026a), but without robustly assessing whether such a value was the optimal one for the different regions monitored.

Despite the large number of user-defined choices involved, no study has systematically evaluated how different species-selection strategies and minimum probability of occurrence thresholds affect BirdNET performance and whether such variations differ among geographic regions, potentially reflecting sampling biases in eBird. Therefore, global-scale research is needed to identify the best criteria for optimizing BirdNET performance. To address this gap, we provide a comprehensive evaluation of BirdNET performance in the context of bird community characterization under three species selection criteria: 1) the full list of species available in BirdNET, 2) location-based lists for the whole year, and 3) spatio-temporally constrained lists generated for specific weeks of the year and locations. Once the optimal species selection criterion was identified, defined as the one optimizing the balance between BirdNET precision and recall—as measured by precision-recall area under the curve (PR AUC) scores—for describing bird communities, we assessed the impact of a wide range of minimum probability of occurrence thresholds (from 0.01 to 0.1). Our study focuses on the ability of BirdNET to describe bird communities, rather than to detect bird vocalizations, since the different species selection criteria evaluated here affect the number of species included in BirdNET output, with no impact on the number of vocalizations detected per species. With this aim, we analyzed 5,047 minutes of audio collected from 72 recording locations across six continents, comprising a total of 1,192 bird species annotated by local expert ornithologists. We expect that our results will guide future studies using BirdNET in selecting appropriate species-selection lists and will support the further refinement of BirdNET-based bird monitoring.

## MATERIAL AND METHODS

### SOUNDSCAPE COLLECTION

We used all the soundscapes from the World Annotated Bird Acoustic Dataset (WABAD, version 3, Pérez-Granados et al. 2026b), which comprised 5,047 minutes of audio annotated at the vocalization level by local experts. Recordings were collected from 72 recording locations across six continents (Figure 1). Although most recording locations are found in Europe and North America, there are also 17 datasets from South America (Colombia, Brazil, and Argentina), four from Africa (Guinea-Bissau, Burkina Faso, Cameroon and Uganda), six from Asia (Indonesia, Taiwan, Vietnam, and mainland China) and five from Oceania (Hawaii, New Caledonia, and New Zealand). Detailed information, including the recording location, minutes annotated for each location, and the main biome, is provided in Supplementary Table S1. The recordings and annotations used in this study are publicly available in Pérez-Granados et al. (2026b).

**Figure 1:**
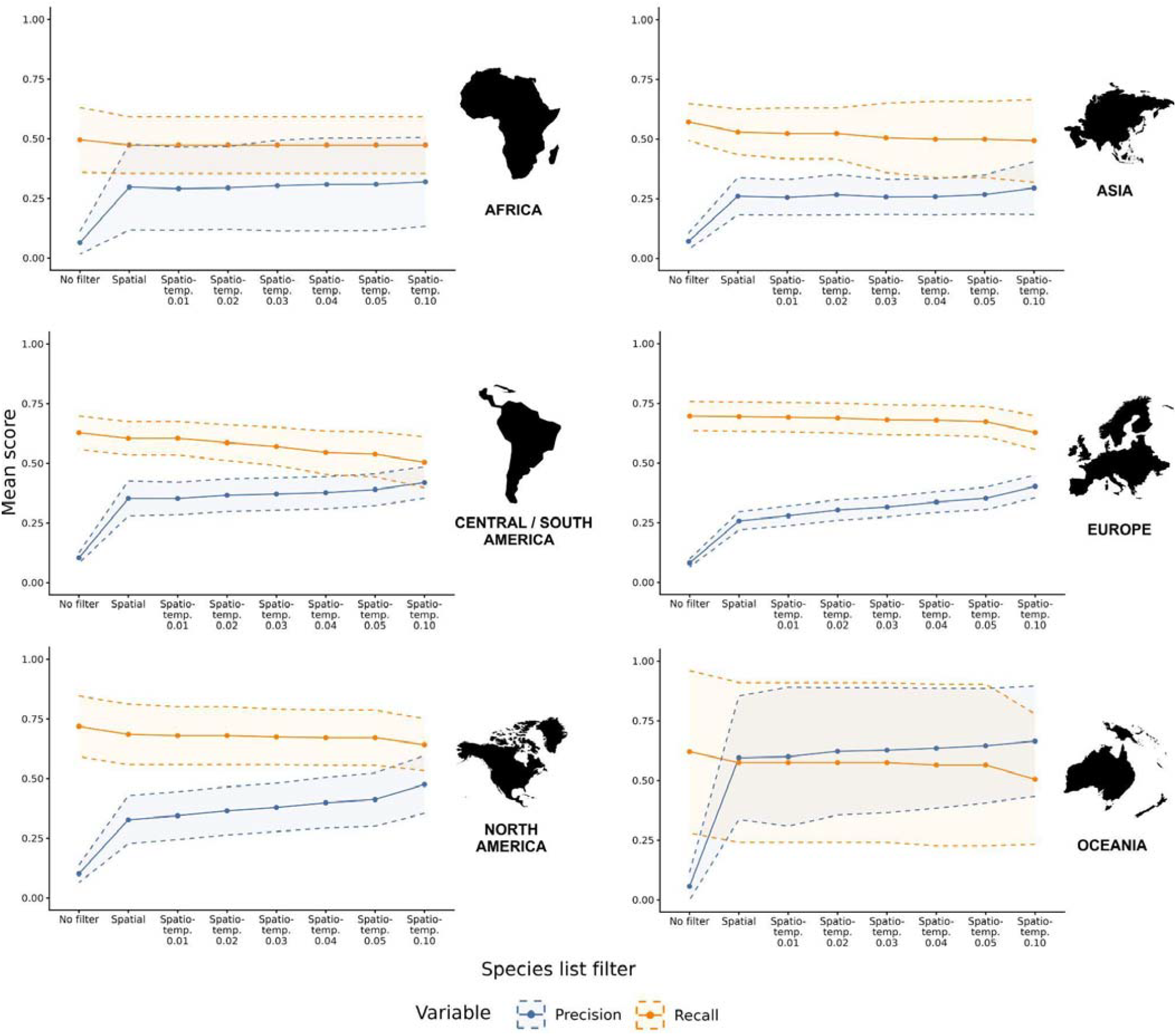
Average precision (blue) and recall (orange) scores, along with 95% confidence intervals, for each species list configuration considered using a minimum confidence threshold of 0.1. Values are shown separately for each world region.

### AUDIO ANNOTATIONS

Recordings in WABAD were annotated by experienced ornithologists with extensive knowledge of the local bird communities. Species names followed the Clements Checklist (Clements et al., 2021), ensuring consistency with the taxonomy adopted by BirdNET. Vocalizations were annotated by drawing bounding boxes on spectrograms of the recordings, where the horizontal limits represent the start and end times of the vocalization and the vertical bars mark their frequency range, from minimum to maximum. Multiple vocalizations of the same bird species could be grouped within a single bounding box when the interval between them was shorter than one second; otherwise, they were annotated as two independent vocalizations. The coordinator of each recording location verified that the annotations complied with the established criteria, and approximately 30% of the recordings per site were additionally reviewed by the WABAD coordinators. Each annotation used in this study was made by a single expert ornithologist, without any formal inter-observer agreement assessment. Further details regarding the annotation workflow and access to the full set of audio annotations are provided in Pérez-Granados et al. (2026b).

### BIRDNET ANALYSES

Recordings were analyzed using BirdNET-Analyzer v2.4.0 via its graphical user interface (GUI). Analyses were run using the default *Minimum Confidence Score* (value of 0.1, range 0.01-1). Based on recent global-scale assessments of BirdNET performance (Pérez-Granados et al. 2026a), we set an *Overlap* of 2 seconds (range 0-3 seconds), a value that maximized BirdNET performance at the dataset level. High *Overlap* values increase the probability that an entire vocalization is contained within a single prediction window, thereby improving species detectability (Pérez-Granados et al. 2026a). *Sensitivity* was set to the default value of 1.0 (range 0.5-1.5) given the high cross-region variability in the value optimizing BirdNET performance for community-level analyses (Pérez-Granados et al. 2026a). Recordings were analyzed with BirdNET, enabling predictions for all sound categories available in BirdNET-Analyzer v2.4 (https://github.com/birdnet-team/BirdNET-Analyzer/tree/main/birdnet_analyzer/labels/V2.4). This configuration corresponds to the first alternative of the three species list configurations considered in the study (“*No filter*”).

The species lists corresponding to the other two configurations were generated via a Linux shell script interfaced with the algorithm’s Python backend, following the approach described in Funosas et al. (2024), and were used to filter BirdNET output obtained with the “*No filter*” configuration. The second species list configuration (“*Spatial filter*”) included all potentially detectable species at each recording site over the entire year, and the third configuration (“*Spatio-temporal filter*”) included the species potentially detectable at each recording site and week of the year. Under this configuration, species lists were generated separately for each week of recording at a given site, such that sites sampled over multiple weeks could have different species lists (i.e., one list per week). For both *Spatial* and *Spatio-temporal* filtering approaches, species lists were extracted using the default minimum probability of occurrence threshold (0.03). After identifying the most appropriate species selection criterion, we further evaluated the effect of the probability of occurrence setting on BirdNET performance by generating additional spatio-temporally filtered species lists using minimum probabilities of occurrence of 0.01, 0.02, 0.03 (default value), 0.04, 0.05, and 0.1.

### ASSESSMENT OF BIRDNET PERFORMANCE

We assessed the performance of BirdNET under the three species list configurations by comparing its predictions with expert annotations using a set of custom scripts implemented in R (version 4.2.2; R Core Team 2025). These scripts were adapted, and are publicly available, from those described in Funosas et al. (2024). Performance was evaluated at the dataset level, reflecting the capacity of BirdNET to characterize bird community composition from multiple recordings collected at a single site. BirdNET predictions were classified into four categories:

- **True Positives (TP):** A species was considered a TP if BirdNET correctly identified that species at least once in any recording from the focal site.
- **False Positives (FP):** A species was classified as a FP when all BirdNET predictions of that species at the focal site were incorrect.
- **True Negatives (TN):** A species was considered a TN when it was neither predicted by the algorithm nor detected by the expert annotator.
- **False Negatives (FN):** A species was classified as a FN when it was identified by the expert but not predicted by BirdNET.

Based on these categories, we evaluated BirdNET precision, recall, and F1-scores at dataset level. Precision represents the proportion of correctly identified species among all species predicted by BirdNET, whereas recall represents the proportion of expertLidentified species correctly detected by the algorithm (Knight et al. 2017, Pérez-Granados 2023). Importantly, recall calculations included species outside BirdNET taxonomic coverage (i.e., species not covered in the most recent BirdNET version). Any annotated species undetected by the algorithm was classified as a FN, regardless of whether the absence resulted from “*Spatial*” or “*Spatio-temporal filtering*”, from the species falling outside BirdNET taxonomic scope, or from a lack of accuracy of BirdNET to correctly predict a species already included in BirdNET. This decision reflects our primary interest: not the recall conditional to the species list used, but the practical performance expected when running BirdNET in a region, i.e., the fraction of species present that the model detects correctly and the extent to which this fraction varies under the different species list configurations.

Finally, F-scores integrate precision and recall into a single performance metric, with the weight attributed to each component being modulated by a β parameter. An F-score with β = 1, or F1-score, gives equal weight to precision and recall, whereas β > 1 emphasizes recall and β < 1 emphasizes precision. Here, we computed F-scores using β = 1 as a standard value to facilitate comparisons with other studies. Precision, recall and F1-scores were computed across 90 confidence score thresholds ranging from 0.10 to 0.99 in increments of 0.01, following the procedure described by Funosas et al. (2024).

The values obtained for precision and recall were used to estimate the Precision-Recall Area Under the Curve (PR AUC; Davis and Goadrich, 2006). The PR curve plots precision against recall for each minimum confidence threshold considered, illustrating the trade-off between these two metrics. PR AUC values (range 0-1) serve as a measure of the predictive power of the algorithm, with higher values indicating greater predictive power (see similar approaches for BirdNET evaluations in Funosas et al. 2024, 2026). This metric integrates precision across the entire recall range, meaning that extending this range —even toward lower recall values— can increase the total area under the curve. Consequently, a PR curve with a broader recall range can have a higher AUC than one with a narrower range, even if the latter maintains higher precision at every overlapping recall level. Less restrictive species list filters are associated with greater variability in recall scores across confidence scores, resulting in a broader range of recall values in comparison to those obtained with more restrictive species list filters. Hence, to ensure comparability across different species list configurations, PR AUC was adjusted for the recall range using the formula described in Funosas et al. (2026):

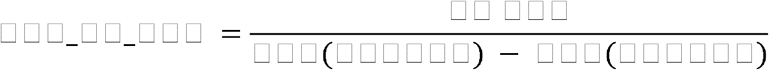

## RESULTS

At the global scale, the number of species correctly and incorrectly predicted, as well as the number of undetected present species, varied strongly across the three species list configurations (Figure 1). BirdNET recall was the highest under the “*No filter*” configuration (0.657), but it resulted in a very low precision (0.186) and the lowest F1-score (0.291, Table 1). Applying a “*Spatial filter”* slightly decreased the recall (0.632), but substantially reduced the number of misclassifications and therefore yielded higher precision (0.445), greatly improving the F1-score with respect to the “*No filter*” approach (0.522, Table 1). BirdNET performance achieved the highest precision (0.471) and F1-score (0.531) when applying a “*Spatio-temporal filter*”, despite a slight reduction in recall (0.609) compared to the other two configurations (Table 1).

**Table 1:**
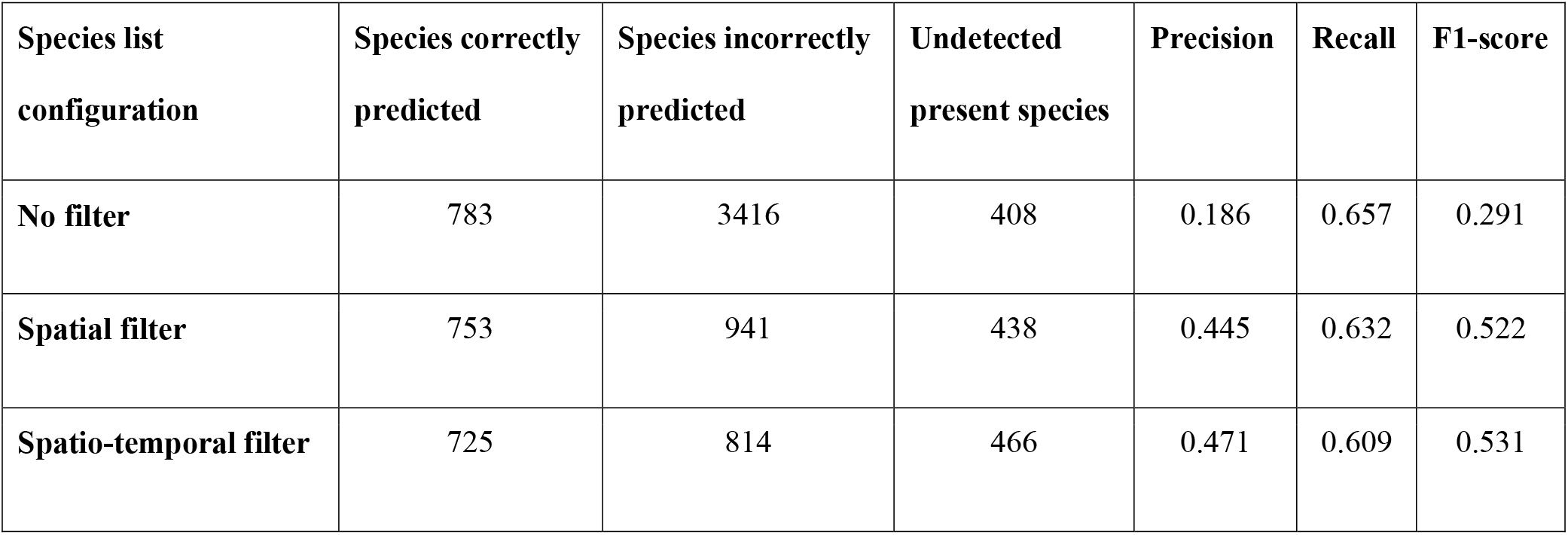
Summary of BirdNET performance on the World Annotated Bird Acoustic Dataset under three species list configurations: 1) considering all species covered by BirdNET (“*No filter”*), 2) applying a “*Spatial filter”* (all species potentially detectable at a given location over the whole year), and 3) applying a “*Spatio-temporal filte*r” (all species potentially detectable at a given location and week of recording). For each configuration, we report the number of species correctly and wrongly predicted by BirdNET, the number of undetected present species, and the precision, recall and F1-score of the algorithm when using a confidence threshold of 0.1.

| Species list configuration | Species correctly predicted | Species incorrectly predicted | Undetected present species | Precision | Recall | F1-score |
| --- | --- | --- | --- | --- | --- | --- |
| No filter | 783 | 3416 | 408 | 0.186 | 0.657 | 0.291 |
| Spatial filter | 753 | 941 | 438 | 0.445 | 0.632 | 0.522 |
| Spatio-temporal filter | 725 | 814 | 466 | 0.471 | 0.609 | 0.531 |

The “*Spatio-temporal filter*” was consistently found to be the optimal species list configuration, based on recall-adjusted PR AUC scores, in all the regions and across most of the datasets analyzed (Table 2). In fact, the “*Spatio-temporal filter*” was identified as the optimal configuration in 61 of the 72 datasets (84.7%) and showed high cross-dataset convergence, both within (range 70–100%) and among regions (Table 2).

**Table 2:** Best species list configuration for each world region as defined in the World Annotated Bird Acoustic Dataset. Three configurations (“*No filte*r”, “*Spatial filter*” and “Spatio-temporal *filter*”) were evaluated based on recall-adjusted PR AUC scores. The number and percentage, relative to the total for each region, of regional datasets converging on the same optimal species list configuration are presented.

| World region | Best species list configuration | Cross-dataset convergence |
| --- | --- | --- |
| **Africa** | Spatio-temporal filter | 3/4 (75%) |
| **Asia** | Spatio-temporal filter | 5/6 (83%) |
| **Central-South America** | Spatio-temporal filter | 14/20 (70%) |
| **Europe** | Spatio-temporal filter | 24/27 (89%) |
| **North America** | Spatio-temporal filter | 10/10 (100%) |
| **Oceania** | Spatio-temporal filter | 5/5 (100%) |

When using the “*Spatio-temporal filter*” configuration, increasing the minimum occurrence frequency threshold from 0.01 to 0.1 led to a progressive increase in precision (from 0.446 to 0.516, respectively), at the cost of a gradual decrease in recall (from 0.630 to 0.559, respectively, Figure 1). As a result, the F1-score remained relatively stable across the six minimum occurrence frequency thresholds analyzed (range 0.522 - 0.539, see Table 3). The 0.05 minimum occurrence frequency threshold provided the best balance between precision and recall for community-level descriptions.

**Table 3:** Summary of BirdNET performance on the World Annotated Bird Acoustic Dataset when using a “*Spatio-temporal filter*” under six minimum occurrence frequency thresholds: 0.01, 0.02, 0.03, 0.04, 0.05, and 0.1. For each threshold, we report the number of species correctly and incorrectly predicted by BirdNET, the number of undetected present species, and the precision, recall, and F1-score of the algorithm when using a confidence score threshold of 0.1.

| Minimum occurrence frequency threshold | Species correctly predicted | Species incorrectly predicted | Undetected present species | Precision | Recall | F1-score |
| --- | --- | --- | --- | --- | --- | --- |
| **0.01** | 750 | 930 | 441 | 0.446 | 0.63 | 0.522 |
| **0.02** | 739 | 859 | 452 | 0.462 | 0.62 | 0.53 |
| **0.03** | 725 | 814 | 466 | 0.471 | 0.609 | 0.531 |
| **0.04** | 711 | 761 | 480 | 0.483 | 0.597 | 0.534 |
| **0.05** | 705 | 722 | 486 | 0.494 | 0.592 | 0.539 |
| **0.1** | 666 | 625 | 525 | 0.516 | 0.559 | 0.537 |

## DISCUSSION

The use of BirdNET for automated bird monitoring has expanded rapidly and continues to increase (Funosas et al. 2026, Mei et al. 2026, Pérez-Granados et al. 2026a, Petry et al. 2026). Here, we analyzed an extensive acoustic dataset annotated by local experts from 72 locations worldwide (Pérez-Granados et al. 2026b) to provide a comprehensive evaluation of BirdNET capacity to characterize bird communities under three different species list configurations. Our results reveal that species filtering implies a trade-off between precision and flexibility for applying automated bird recognition at large spatial scales. The use of both “*Spatial*” and “*Spatio-temporal filters*” substantially improved BirdNET precision over the “*No filter*” configuration, but at the cost of a slight reduction in the number of correctly predicted species. However, since the incorrect species predictions filtered out greatly exceeded the loss of correctly predicted species, overall BirdNET performance improved significantly when using species filters. In particular, “*Spatio-temporal filtering*” was identified as the optimal strategy, maximizing BirdNET performance in most datasets and consistently across the six world regions analyzed. We also identified that intermediate minimum occurrence thresholds (0.05) maximized BirdNET performance (as measured by F1-scores), as the use of higher thresholds reduced recall more than they increased precision, and vice versa for lower thresholds.

Although our study focuses on the effects of different filtering approaches on the potential of BirdNET to characterize bird communities, the applicability of our results extends to other PAM applications. For example, using filters in studies aiming to monitor avian phenology from sound recordings (Bota et al. 2020, Van Doren et al. 2023, Sethi et al. 2024) may prevent the detection of early and late migrants in the dataset, when the species’ probability occurrence is expected to be very low. Likewise, species characterized by low abundance, low detectability or that are difficult to identify, and therefore likely underrepresented in citizen-science programs such as eBird, may also be excluded from BirdNET autogenerated species lists or remain poorly represented. In such cases, mismatches between true and expected occurrences may lead to their exclusion from species lists for particular locations or time periods. Moreover, the use of such filters may hinder the detection of threatened birds in newly surveyed locations (Mosikidi et al. 2023). In many of these cases, replacing automated filtering approaches with custom-defined species lists tailored to the target species communities may be beneficial. Overall, although our results provide guidance on the optimal filtering approaches for future BirdNET surveys, the selection of such configuration and thresholds should be carefully considered in light of the specific ecological questions being addressed.

It is also important to acknowledge that, even if the use of species filters improves output interpretability and reduces false positives, it also inherently reduces the number of predictions provided by the algorithm. In contrast, an unfiltered approach preserves the full range of model predictions, allowing experts to manually verify and reclassify predictions at a posterior stage. This might be useful in certain research contexts, since most of the discarded predictions are actual bird vocalizations. A potential direction for future BirdNET updates to deal with that issue would be to incorporate biogeographic plausibility into automated species classification. At present, predictions corresponding to species not occurring at a given recording location are removed when “*Spatial filtering*” is applied. However, BirdNET could explore options to reclassify model predictions originally assigned to species not occurring locally to closely related or acoustically similar species that do occur in the recording location. We acknowledge that implementing such functionality might be challenging and extend beyond both the scope of standard automated classification, but it could help reduce the risk of discarding detected bird vocalizations while maintaining spatial realism and facilitating posterior validation.

Despite the broad spatial scale considered in our study, a substantial majority of the acoustic datasets analyzed come from Europe (27 datasets) and the Americas (30 datasets), with Africa, Asia, and Oceania sparsely represented (4-6 datasets each, Table 2). This imbalance among regions is particularly relevant because species lists generated in BirdNET-Analyzer are expected to be less reliable in regions with limited eBird data coverage, such as Africa and Asia. The scarcity of acoustic datasets from these regions in our study constrains our ability to provide a robust global assessment of the impact of the different species list configurations on BirdNET performance. Recent research has highlighted that BirdNET performance is poorer in world regions with lower eBird data coverage relative to well-sampled regions (i.e., higher BirdNET performance in Europe and the Americas, Funosas et al. 2026). The lower performance of BirdNET in these areas might be related to the lower amount of training data to develop robust classifiers, but a potential, non-exclusive, explanation is that the inaccuracies in the automatically generated species lists may further exacerbate performance limitations in such areas. Therefore, although our results are consistent within and among regions, they should be interpreted with caution in regions with limited data availability, and additional assessments should be conducted to better understand the impact of species list configurations on region-specific BirdNET performance.

In this study, we provide the first assessment of the impact of species list configurations on BirdNET performance. Our broad spatial scope, together with the consistent improvement observed when using a “*Spatio-temporal filtering*” approach, suggests that this strategy can enhance the robustness of future bird surveys using BirdNET. These results may also inform future updates of BirdNET, for example by improving the documentation on the effects of species list configuration settings or by refining the minimum occurrence threshold default value to 0.05 for community-level analyses. We recommend that users verify BirdNET-generated species lists before using them as filters, particularly in contexts where these lists may strongly influence results, such as phenology studies, rare or vagrant species detection, or regions with limited eBird data coverage. We suggest that this verification step should be incorporated into standard BirdNET workflows. Our assessment also highlights the importance of expanding the availability, accessibility, and adoption of biodiversity monitoring platforms, such as eBird, in underrepresented regions, as enhanced data coverage would directly enhance occurrence models and methodological development. Likewise, increasing the availability of field recordings in these regions, as well as the publication of curated annotated acoustic datasets such as the one used here (Pérez-Granados et al. 2026b), may also support further refinement of BirdNET and other machine learning tools (Morfi et al. 2019, Ghani et al. 2023).

## ACKNOWLEDGEMENTS

CPG was funded by the Ramón y Cajal 2024 Programme (RYC2024-048830-I) of the Spanish Ministry of Science, Innovation and Universities, funded by MICIU/AEI/10.13039/501100011033 and FSE+. JM was funded by the Basque Government Postdoctoral grant (POS_2025_1_0009). ESG received the grants RYC2019-027216-I and CNS2023-144791 funded by the Spanish Ministry of Science, Innovation and Universities (MCIN/AEI/ 10.13039/501100011033) and by ESF Investing in your future.

